# Biochemical assessment of α-α-subunit interactions of Na_v_1.5 in a heterologous expression system

**DOI:** 10.1101/2025.10.02.679760

**Authors:** Oksana Iamshanova, Anne-Flore Hämmerli, Suberja Sundaralingam, Arbresh Seljmani, Sabrina Guichard, Maria Essers, Jean-Sébastien Rougier, Hugues Abriel

**Author notes:** Correspondence should be addressed either to: Oksana Iamshanova, PhD or Abriel Hugues, MD, PhD, Institute of Biochemistry and Molecular Medicine, University of Bern Bühlstrasse 28, CH-3012 Bern, Switzerland.

## Abstract

Heterologous overexpression of any protein, and especially of the large transmembrane channel Na_v_1.5, could be associated with the insufficiency of endoplasmic reticulum folding machinery, hence leading to aspecific protein aggregation indistinguishable from the genuine α-α-subunit interactions. In this study, we show that the interactions between heterologous Na_v_1.5 proteins depend on nascent N-linked glycosylation, are supported by non-native intermolecular disulfide bonds, and are likely predisposed to hydrophobic “stickiness”. Particularly, we show strong interactions between the full-length Na_v_1.5 and its truncated peptides: N-terminal domain, all four transmembrane domains, as well as the intracellular linker between domains I and II. Taken together, we conclude that the heterologous expression system is not optimal for the identification of α-α-subunit interaction sites of Na_v_1.5, and this question needs to be further addressed in the native tissues.

**Graphical abstract:** 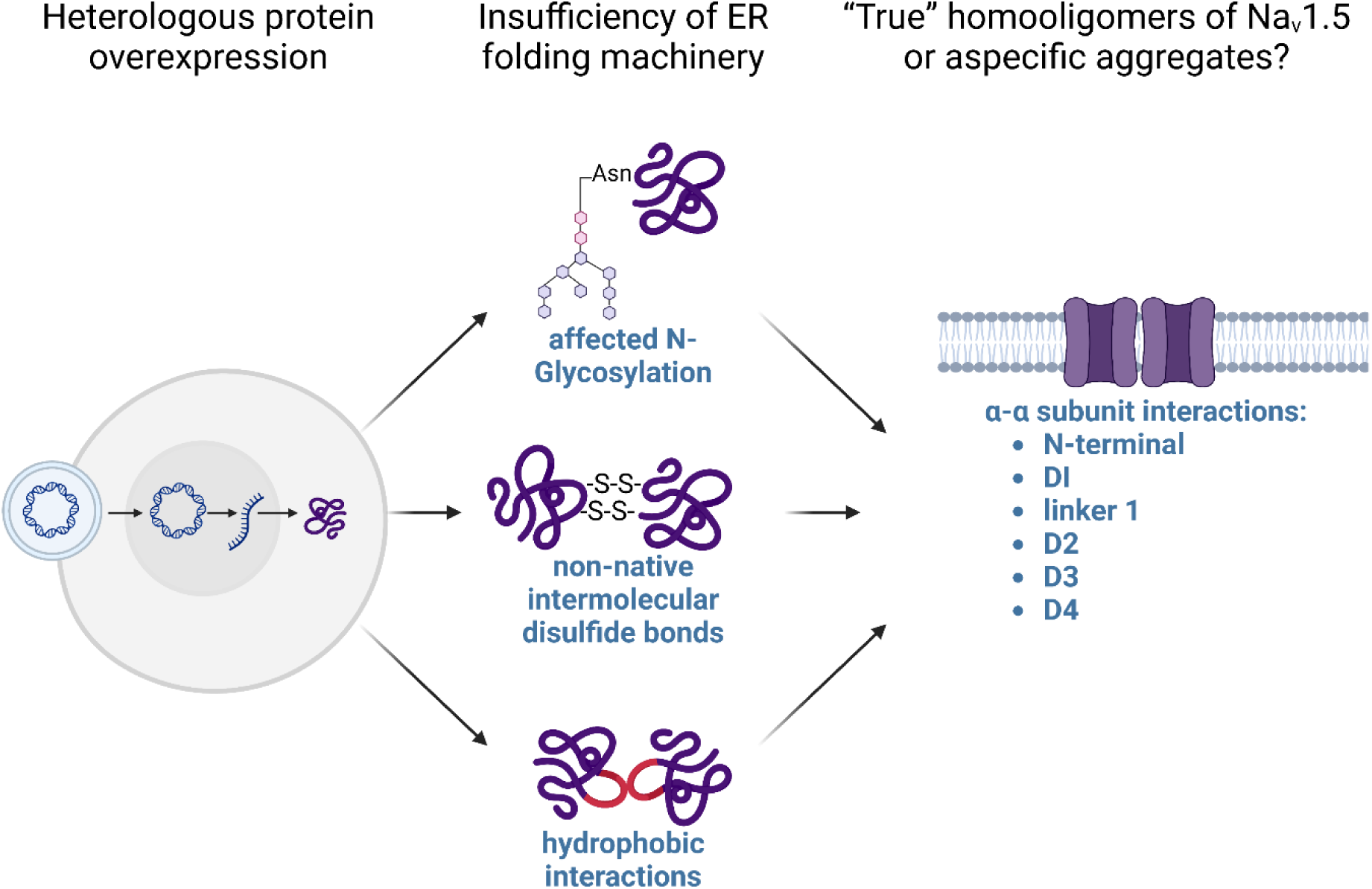

## Introduction

Human voltage-gated sodium (Na^+^) channels (VGSCs) represent a family of proteins consisting of one pore-forming α-subunit (Na_v_1.x) and two distinct auxiliary β-subunits (Na_v_β) (Shen et al., 2019) The pore-forming α-subunit is comprised of four transmembrane domains (I–IV), each of which has six transmembrane segments (S1– S6), where S1–S4 represent the voltage-sensor and S5 and S6 form the pore (Li et al., 2021). N- and C-terminals, as well as linkers between the transmembrane domain (hereafter named L1, L2, and L3) are intracellular (Li et al., 2021). The most distinguishable feature of VGSCs is their fast permeation of Na^+^ current (*I*_Na_) in response to membrane depolarizations of few millivolts (mV), followed by rapid inactivation. Therefore, VGSCs are first in line to respond to an initial depolarization and, hence, are the key regulators of cellular excitability. Within various tissues and cell types (e.g., neurons, neuromuscular junctions, and cardiomyocytes), VGSCs have been observed to form clusters (Eshed-Eisenbach and Peles, 2021). The clustering of VGSCs is believed to enlarge their functional output, i.e., increase membrane responsiveness to electrical stimuli and conduction (Dixon et al., 2022; Eshed-Eisenbach and Peles, 2021). However, it remains unclear whether the clustering of VGSCs is supported by direct interactions between its subunits.

While Na_v_β-subunits were shown to homo- and hetero-oligomerize (Bouza and Isom, 2017), α-subunits were thought to present as singular protein units clustered together, mainly due to specific membrane targeting (Marchal and Remme, 2022). Not long ago, this dogma was challenged by findings reporting direct α–α-subunit interactions. Specifically, human Na_v_1.1, Na_v_1.2, Na_v_1.5, and Na_v_1.7 were shown to homodimerize when heterologously expressed without Na_v_β-subunits (Clatot et al., 2012, 2017; Rühlmann et al., 2020; Mercier et al., 2012; Zheng et al., 2021; Iamshanova et al., 2024). Importantly, the bacterial α-subunit of VGSC was previously demonstrated to assemble as a dimer-of-dimers, consequently forming a functional ion-conducting homotetrameric channel (Payandeh et al., 2011). Domain-swapping tetramerization of the bacterial α-subunit of VGSC was reportedly achieved by hydrophobic, polar, and charged amino acid residues mainly located within the S5 and S6 helices (Payandeh and Minor, 2015). Higher-order oligomers were not observed for the prokaryotic α-subunit of VGSC until a recent study suggested inter-channel dimerization may occur through the voltage-sensing domain in the resting state (Sumino et al., 2023). In contrast, human Na_v_1.5 reportedly dimerizes through its intracellular L1 region between domains I and II (Clatot et al., 2017). L1 is a subject of multiple post-translational modifications, in particular, the α–α-subunit interaction region described for Na_v_1.5 (i.e., Arg493-Arg517) is comprised of at least one methylation site and several phosphorylation sites (Marionneau and Abriel, 2015). However, nothing is known yet about the dependence of Na_v_1.5 dimerization on its activation state or relationship to post-translational modifications.

Biosynthesis of Na_v_1.5 is a multistep fine-tuned process starting with *SCN5A* transcription, followed by the production of nascent peptide in the endoplasmic reticulum (ER), its maturation, and subsequent anchoring to the plasma membrane (Dong et al., 2020). Typically, faulty and nonfunctional channels are trapped in the ER, triggering the unfolded protein response (UPR) that leads to the degradation of unwanted protein cargo (Wiseman et al., 2022). UPR is especially important for adult cardiomyocytes, where native Na_v_1.5 is abundant, to maintain cellular functionality because their regenerative potential is significantly low (Liu and Dudley, 2018). Folding of Na_v_1.5 is regulated by the ER quality control machinery, comprised of chaperones and folding enzymes (Dong et al., 2020). As the nascent polypeptide chain enters the ER, a core glycan is added to an Asn at a specific N-glycosylation site (Wiseman et al., 2022). Monoglucosylated N-linked glycans are substrates for the lectin chaperones calnexin and calreticulin, which assist in oxidative folding (Wiseman et al., 2022). Another family of ER-residing folding enzymes is the protein disulfide isomerase family, members of which catalyze disulfide formation, isomerization, or reduction between juxtaposed Cys (Wiseman et al., 2022). Thus, rapid detection and elimination of unfolded nascent peptides in the ER are complicated due to their shared properties with folded unmature proteins, such as exposed hydrophobic patches or aggregation-prone regions (Wiseman et al., 2022). Accordingly, accumulated misfolded proteins can aggregate into larger structures, leading to ER stress and UPR activation (Wiseman et al., 2022). Given the complexity of membrane protein folding, one can hypothesize that in a heterologous expression system, where the host ER machinery may not be fully tailored for proper Na_v_1.5 processing, at least some portion of overexpressed channels may aggregate. Importantly, at this point, misfolded Na_v_1.5 aggregates are difficult to distinguish from true Na_v_1.5 oligomers because they also contain direct α-α-subunit interactions.

Taken together, the aim of our study was to critically assess α-α-subunit interactions of Na_v_1.5 in a heterologous expression system, striving for a better understanding of the VGSC dimerization phenomenon.

## Experimental procedures

### cDNA constructs

Template cDNA constructs and their derivatives are listed in Table 1.

**Table 1.**
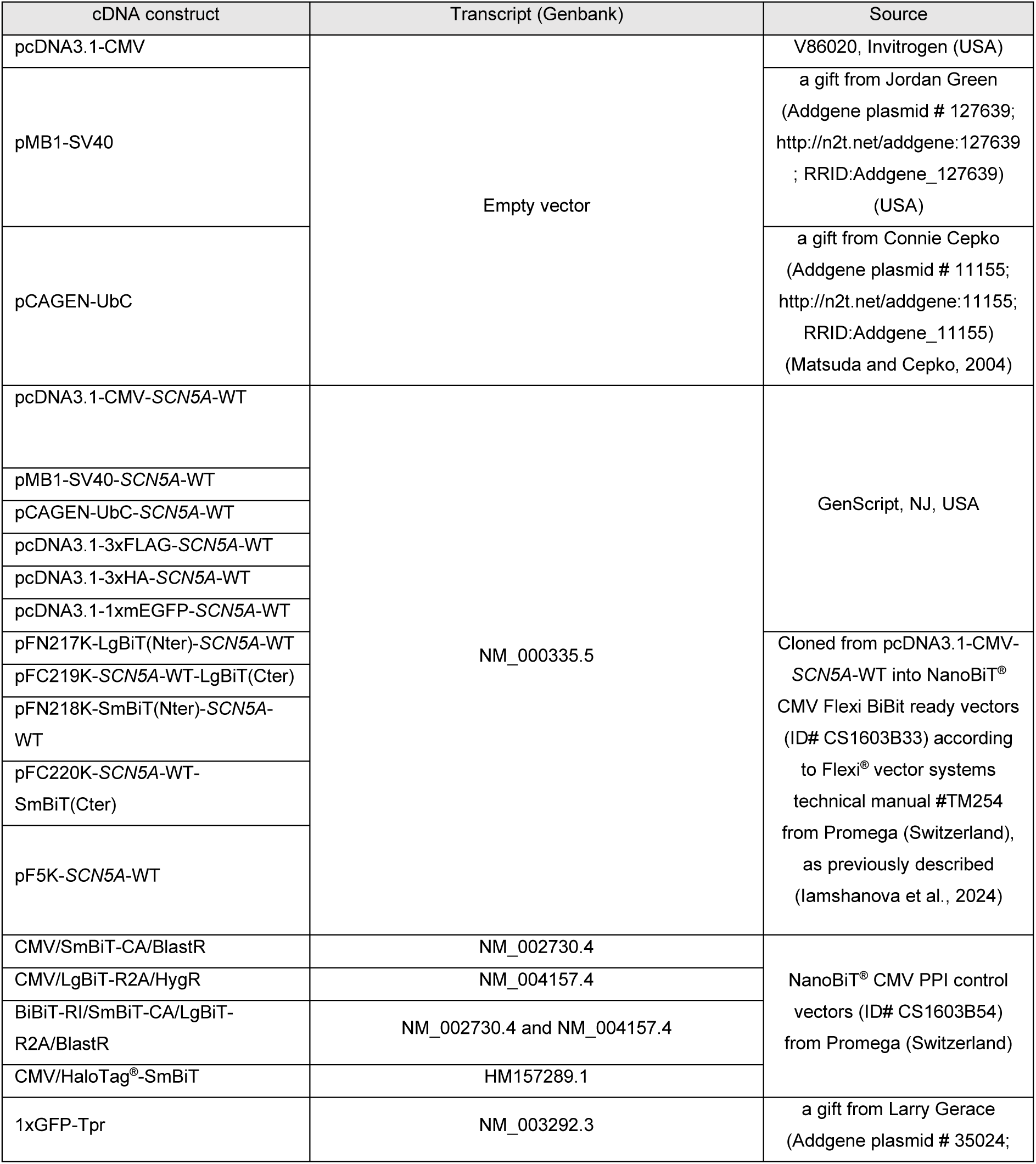
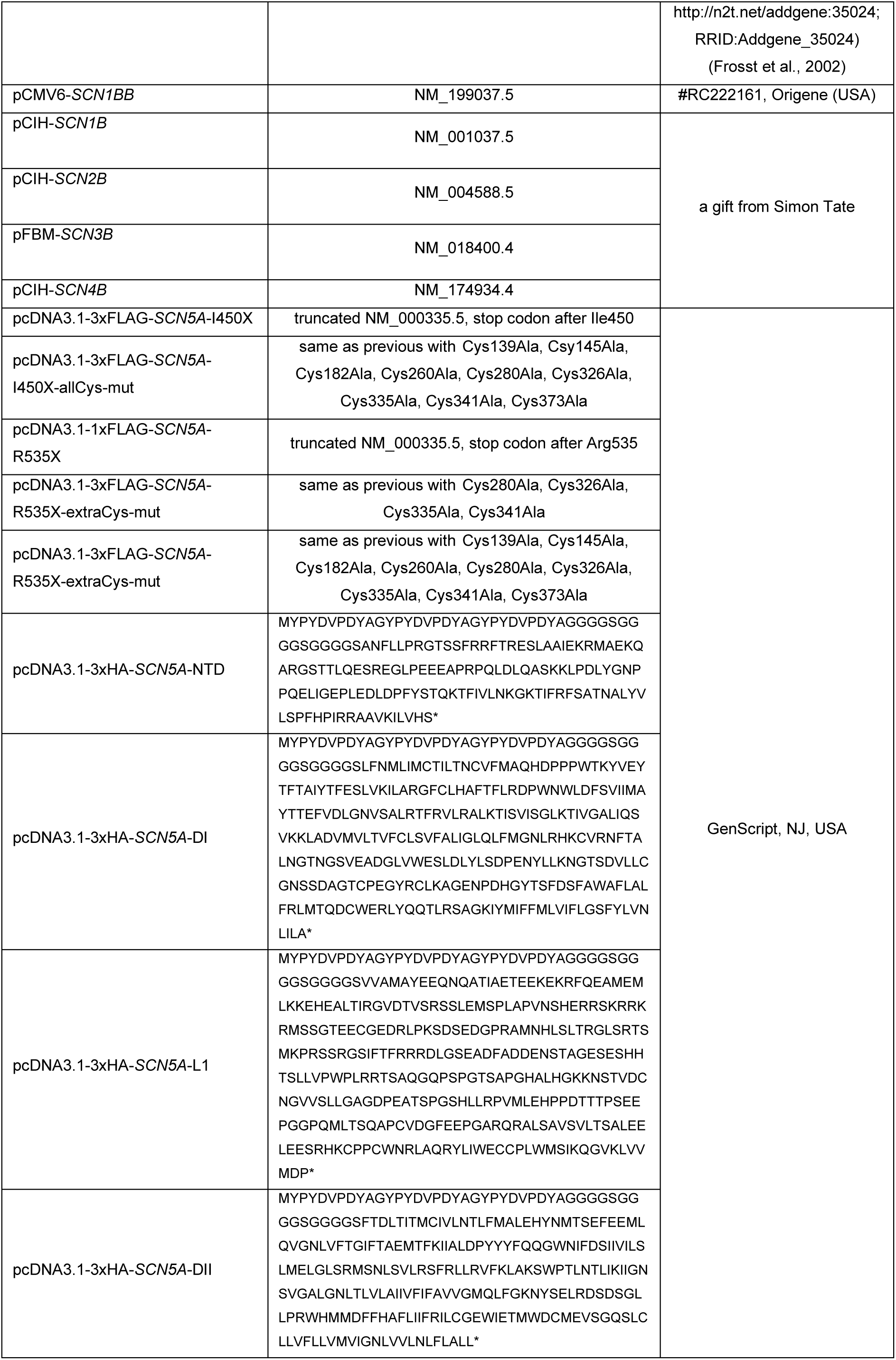
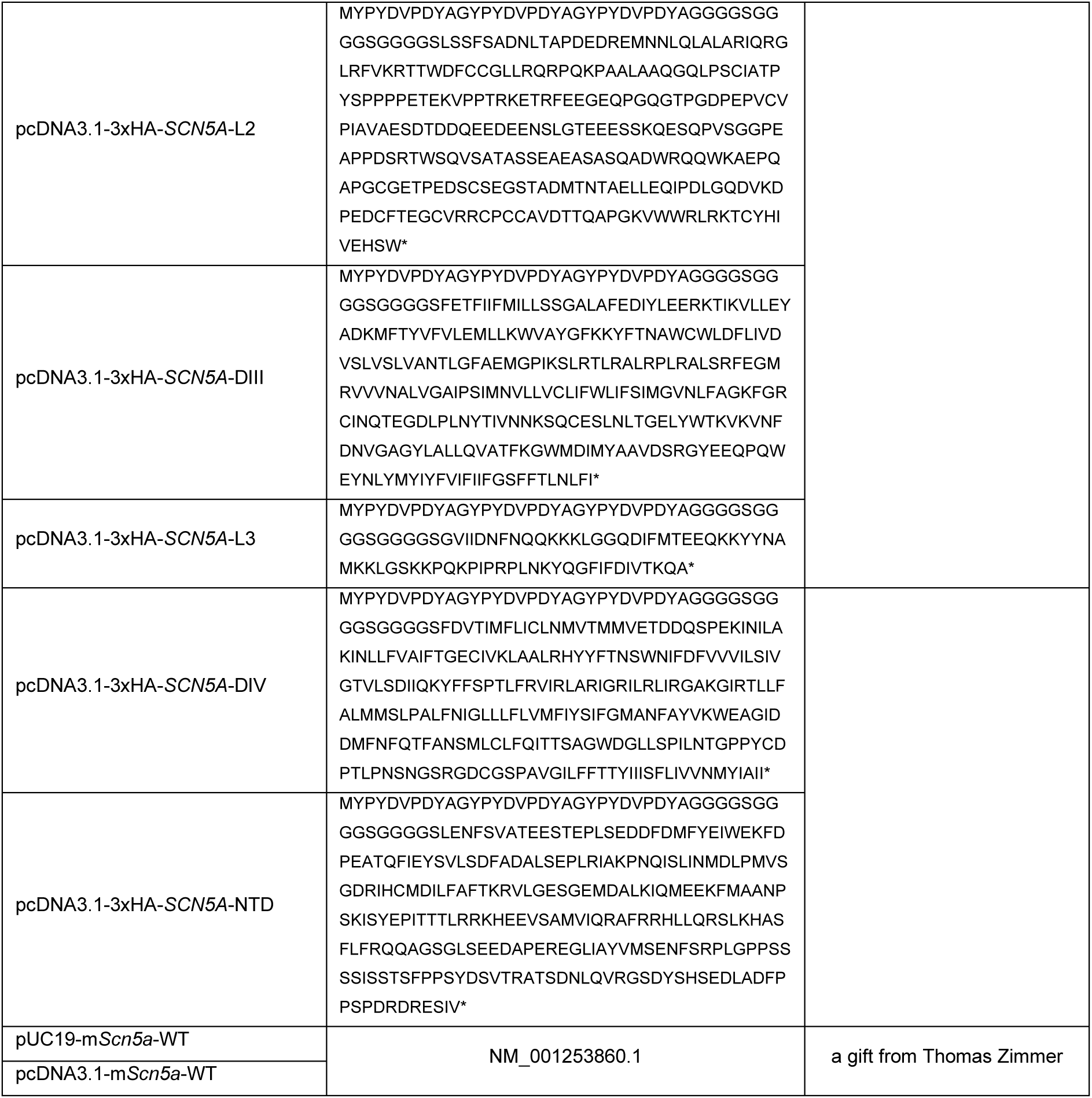
Description of cDNA constructs used in this study.

### Cell culture and transfection

The human embryonic kidney cell line tsA201 (ECACC Cat# 96121229, RRID:CVCL_2737) and monkey kidney cell line COS-7 (ECACC Cat# 87021302, RRID:CVCL_0224) were cultured up to 20 passages at 37°C with 5% CO_2_ in Dulbecco’s Modified Eagle’s Medium (41965, Gibco^TM^, Thermo Fisher Scientific, USA), supplemented with 2 mM L-glutamine (G7513, Sigma, USA), 50 U/mL penicillin-streptomycin (15140122, Gibco) and 10% heat-inactivated fetal bovine serum (10270-106, Lot 2440045, Gibco). When needed, cells were split using phosphate-buffered saline (PBS; 10010-015, Gibco) and 0.05% Trypsin-EDTA (25300-054, Gibco). Mycoplasma contamination status was tested weekly with a PCR Mycoplasma Test Kit I/C (PK-CA91-1096, Promokine, PromoCell GmbH, Germany). cDNA constructs were introduced into cells by transfection with LipoD293^TM^ (SL100668, SignaGen^®^ Laboratories, USA). Cells were harvested 48 hours after transfection unless stated otherwise.

### Protein extraction from cellular monolayers

Pre-washed (PBS) adherent monolayers of cells were scraped in 2 mL of cold PBS (pH 7.4) and pelleted by centrifugation for 5 minutes at 200 × g at 4°C. Cell pellets were lysed in lysis buffer (50 mM NaCl, 50 mM imidazole/HCl, 2 mM 6-aminohexanoic acid, 1 mM EDTA, pH 7) with the addition of cOmplete tablets EDTA-free (04693132001, Roche, Switzerland), 0.5 mM Na_3_VO_4_, 0.5 mM NaFl, 10 μg/mL aprotinin, 10 μg/mL leupeptin, 1 mM phenylmethylsulfonyl fluoride, and 1% digitonin (D141, Sigma) for 1 hour at 4°C. Lysates were centrifuged at 16,100 × g for 15 minutes at 4°C. The obtained supernatant was taken for further analysis. Protein concentrations were determined by Coo Assay Protein Dosage Reagent (UPF86420, Uptima, Israel).

### Protein extraction from mouse heart

According to the Swiss Federal Animal Protection Law, all animal experiments were performed and approved by Bern’s Cantonal Veterinary Administration (license BE88 2022). This investigation conforms to the Guideline for the Care and Use of Laboratory Animals, published by the US National Institutes of Health (NIH publication no. 85-23, revised 1996). Mice were housed in a controlled, specific pathogen-free environment (23 ± 1°C; humidity 60%; lights on 06:00 AM – 06:00 PM; food and water available ad libitum; enriched environment) with a maximum of 6 mice per cage.

Female and male C57BL/6JRj mice between 20-40 weeks old were deeply anesthetized using ketamine/xylazine injection (intraperitoneal; one time; 200/20 mg/kg body weight) with heparin 25000IE/5 ml (4 ml/kg body weight) to avoid blood coagulation. Mice were sacrificed by a thoracotomy (median sternotomy) to access the heart as quickly as possible. Then the heart was rapidly excised and taken for further experimental procedures.

For the isolation of cardiomyocytes, the excised heart was cannulated and mounted in a Langendorff system for retrograde perfusion at 37°C. The heart was perfused free of blood (few minutes) with nominally Ca^2+^-free solution containing (mmol/L): 135 NaCl, 4 KCl, 1.2 MgCl_2_, 1.2 NaH_2_PO4, 10 HEPES, 11 glucose, pH 7.4 (NaOH). Next, the heart was perfused with the same solutions with 50 µM Ca^2+^ and 275U collagenase type II (275U/mg, CLS-2, Worthington, NJ, USA) till digestion. Following digestion, the ventricle was transferred to the aforementioned buffer with 100 µM Ca^2+^, minced into small pieces and triturate to liberate single ventricular myocytes by gentle pipetting, and filtered through a 100 µm nylon mesh. After supernatant was removed, cardiomyocytes were lysed in 1x lysis buffer, 1% digitonin for 1 hour at 4°C.

For the homogenization procedures, the excised heart was washed three times in five volumes of ice-cold PBS and put in one volume of 2x lysis buffer. For homogenization with Benchtop Bioprep-24 (Hangzhou Allsheng Instruments, China), 30 silica beads of 1.4 mm diameter (531607, Milian) and 10 silica beads of 2.8 mm diameter (531608, Milian) were added. The homogenization was performed in 6 cycles at 4260 rpm for 5 seconds (7m/s linear speed) with an interruption of 30 seconds at 4°C between the cycles. For homogenization with VD12 Polytron (VWR), the heart was subjected to 2 cycles of 5 seconds of homogenization on ice.

After homogenization, digitonin was added to make a final concentration of 1% and samples were left to lyse for 1 hour at 4°C. Lysates were centrifuged at 16,100 × g for 15 minutes at 4°C. The obtained supernatant was taken for further analysis. Protein concentrations were determined by Coo Assay Protein Dosage Reagent (UPF86420, Uptima).

### Co-immunoprecipitation

Anti-FLAG^®^ M2 Magnetic Beads (M8823, Sigma) and Pierce^TM^ anti-HA Magnetic Beads (88836, Thermo Scientific) were used at a proportion of 1 μL of beads per 40 μg of total protein lysate. Protein lysates were diluted in Tris-buffered saline [TBS, 20 M Tris, 0.1 mM NaCl, pH 7.6 (HCl-adjusted)] and mixed with 0.05% TBS-Tween-20-prewashed antibody-coupled beads at 4°C overnight. After thoroughly washing the beads with TBS, the co-immunoprecipitated protein complex was eluted with 4× LDS sample buffer (NP0007, Invitrogen, USA). Samples were then analyzed with an immunoblotting technique (see below).

### Cell surface biotinylation

Adherent cell monolayers were gently washed three times with cold PBS (pH 8) and incubated, while slowly rocking, with 1 mg/mL EZlink™ Sulfo-NHS-SS-Biotin (21331, Thermo Fisher Scientific) in PBS (pH 8) for 45 minutes at 4°C. Afterward, excess biotin was quenched by washing with 50 mM Tris-HCl (pH 8), followed by several washes with PBS (pH 7.4). Extracted protein lysates were mixed with Streptavidin Sepharose High-Performance Beads (GE Healthcare, USA) at 4°C overnight. After thoroughly washing the beads with washing buffer [0.1% bovine serum albumin (BSA), 0.001% Tween-20, PBS, pH 7.4], the biotinylated fraction was eluted with 4× LDS sample buffer and analyzed with an immunoblotting technique (see below).

### Sodium dodecyl sulfate–polyacrylamide gel electrophoresis (SDS-PAGE) and immunoblotting analysis

Protein lysates containing LDS sample buffer with or without 100 mM dithiothreitol (DTT) were loaded onto 4%–8% or 4%–12% Tris-acetate acrylamide gels and run at 60V in Tris-acetate running buffer (50 mM Tris, 50 mM Tricine, 0.1% SDS, pH 8.3). Afterward, proteins were transferred to a nitrocellulose membrane using a Trans-Blot Turbo system (1704158, Bio-Rad, USA). After validation of successful protein transfer with Ponceau S solution (0.1% Ponceau S, 12.5% acetic acid), membranes were blocked with 5% BSA in TBS-T (TBS, 0.1% Tween-20) for 1 hour at room temperature and then incubated with primary antibody dilutions at 4°C overnight (Table 1). After incubating membranes with secondary antibodies for 1 hour at room temperature, infrared fluorescent signals were revealed with a LI-COR Odyssey Infrared Imaging System (LI-COR Biosciences, USA) or FUSION FX7 Imaging System (Witec, Germany) and quantified with ImageJ software (Rasband, W.S., US National Institutes of Health, Bethesda, MD, USA). Primary and secondary antibodies used in this study are listed in Table 2.

**Table 2.**
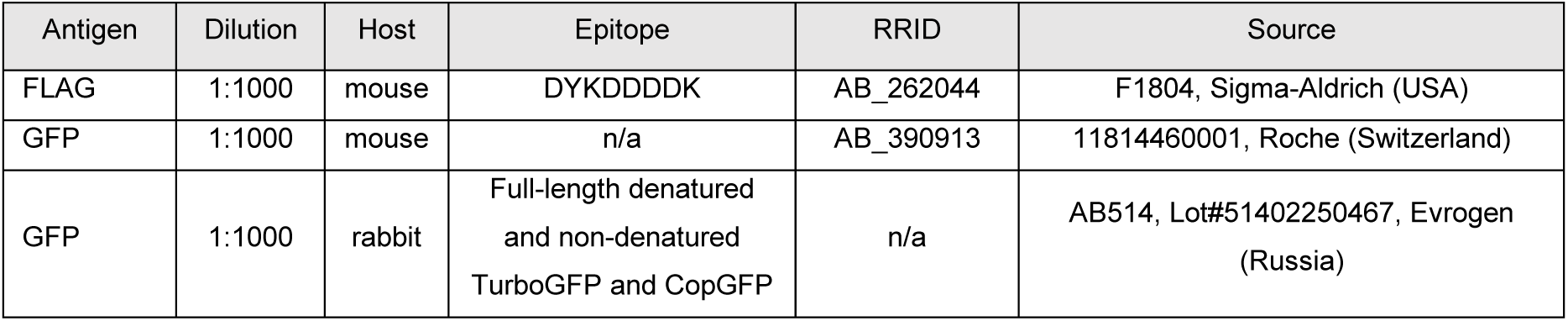
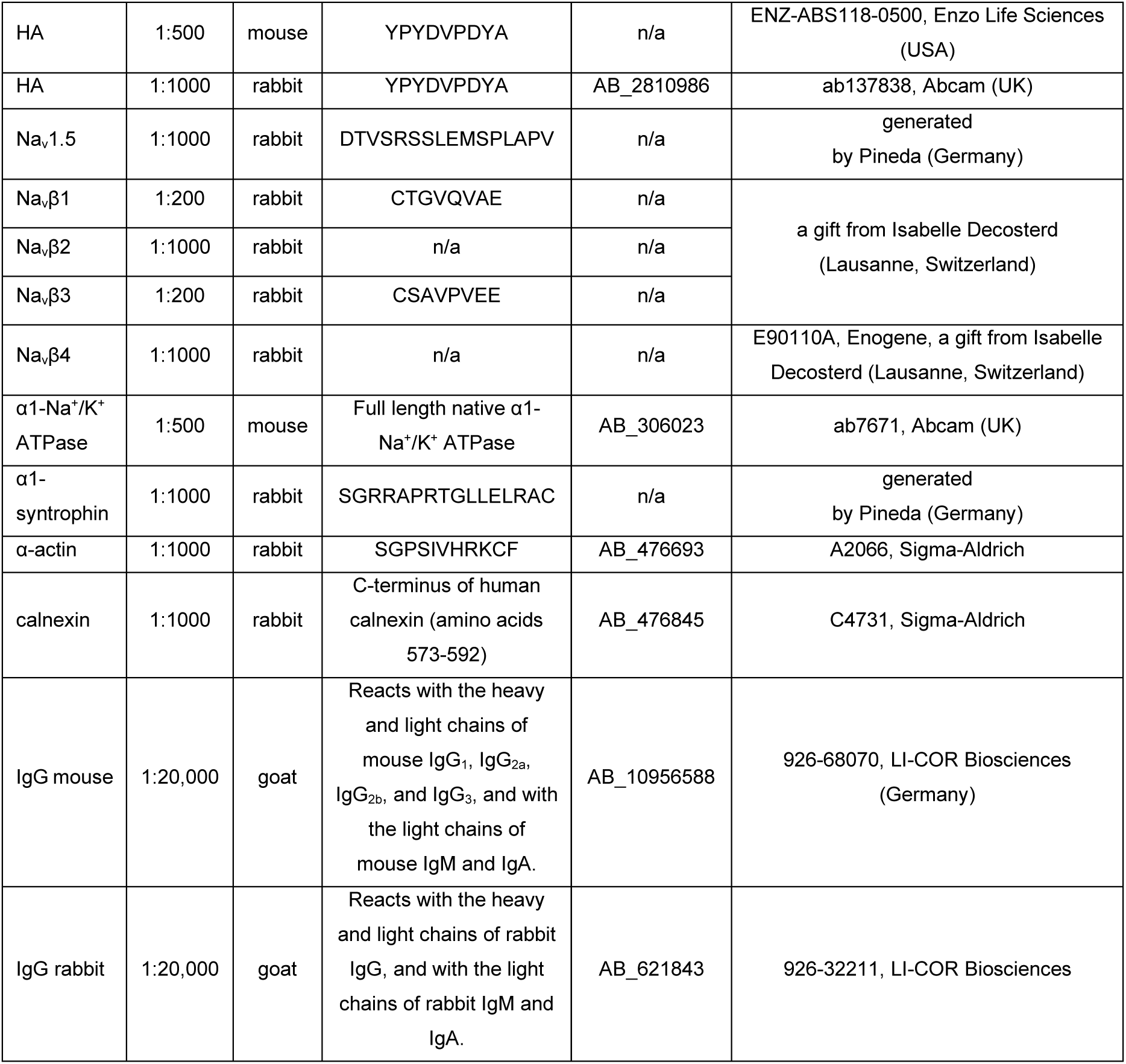
Description of primary and secondary antibodies used in this study.

### Protein-protein interaction assay in living cells

The NanoLuc^®^ Binary Technology NanoBiT^®^ (Promega, USA) allowed us to monitor protein-protein interactions between two Na_v_1.5-WT α-subunits in living cells as described previously (Iamshanova et al., 2024). The luminescent signal was obtained with a Nano-Glo^®^ Live Cell Assay (N2012, Promega) and normalized to the cell quantity represented by the fluorescent signal obtained with a CellTiter-Fluor^®^ Cell Viability Assay (G6082, Promega). Signals were detected with a GloMax^®^ Explorer Multimode Microplate Reader (GM3500, Promega).

### Data and statistical analysis

Data are represented as the mean ± SEM or SD. Data normality was tested using the Shapiro-Wilk test. Statistical significances of normally distributed data were calculated with ordinary one-way ANOVA and Tukey’s multiple comparisons tests. In cases where data were normalized to a control condition, the statistical significance was calculated with a one-sample two-tailed t-test with hypothetical mean value = 1 (Prism version 8.4.3; GraphPad, CA, USA). Exact *p*-values are indicated in the figures. Neither randomization nor blinding were performed in this study.

## Results

### Unlike in native tissue extracts, heterologous expression of NaV1.5 introduces α-α-subunit interactions

Previous studies reported the α-α-subunit interactions of Na_v_1.5 (Clatot et al., 2017, 2012; Iamshanova et al., 2024). It is important to note that the “true” Na_v_1.5 dimers/oligomers and the aggregates of Na_v_1.5 would both possess α-α-subunit interactions. Therefore, hereafter, we use the definition of oligomer alongside protein aggregate, as those two are indistinguishable by most biochemical methods.

Here, we confirmed that differently tagged Na_v_1.5 proteins co-immunoprecipitated in a heterologous expression system represented by tsA201 cells (Fig. 1A). In living cells, the dimerization of Na_v_1.5 was preserved (Iamshanova et al., 2024). Because interactions through the N-N termini of Na_v_1.5 consistently produced the brightest signal (Iamshanova et al., 2024), they were used as a default configuration for the NanoBiT assay in this study. Based on results showing that untagged Na_v_1.5 competed for its interaction with the luminescence-producing pair of Na_v_1.5-LgBiT and Na_v_1.5-SmBiT (Fig. 1B, C) but not the control luminescence-producing pair of well-known interacting proteins PRKAR2A-LgBiT and PRKACA-SmBiT (Fig. S1), we concluded that the observed dimerization was specific to Na_v_1.5.

**Figure 1.**
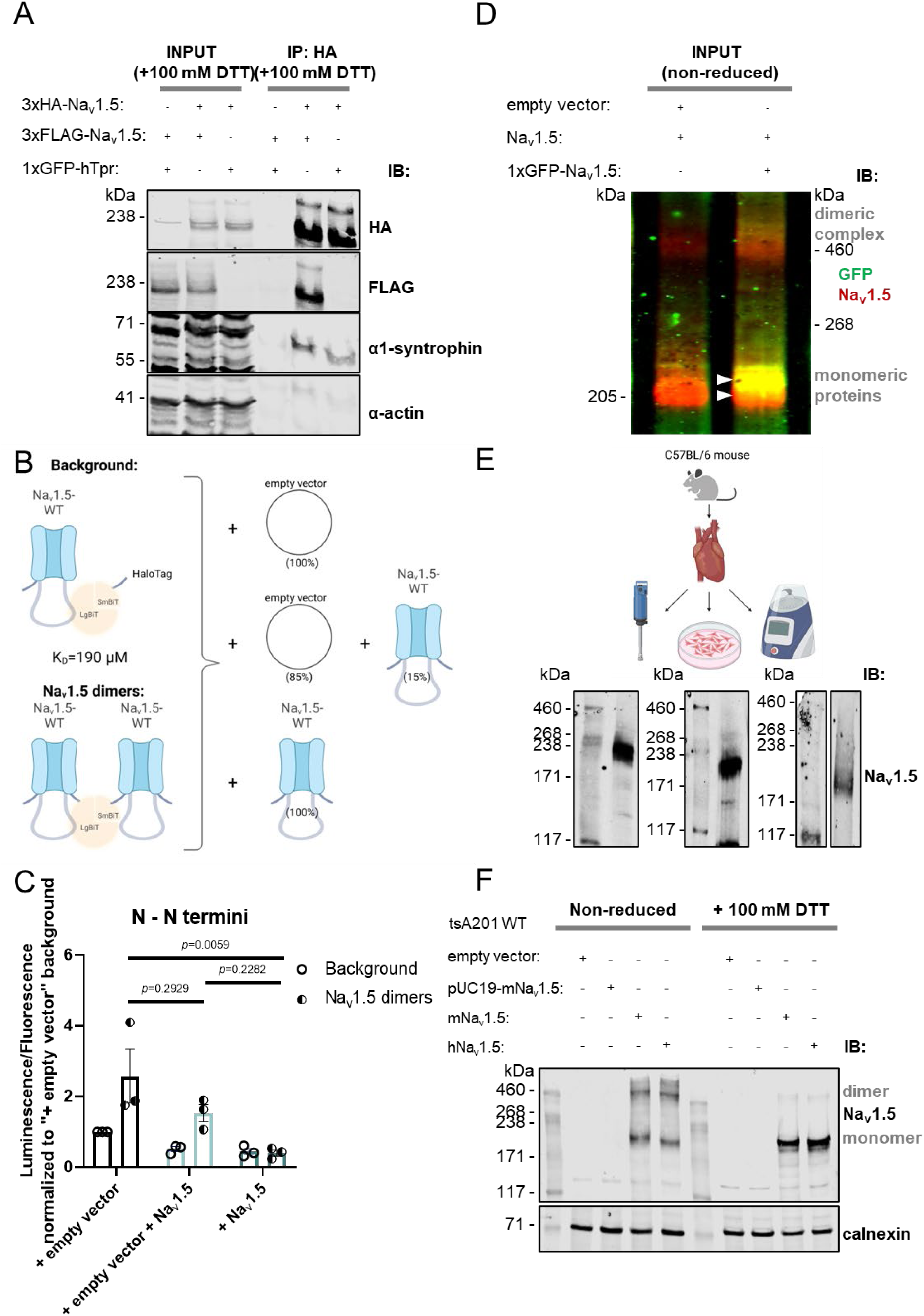
In a heterologous expression system, Na_v_1.5 proteins co-immunoprecipitate and interact in living cells. Na_v_1.5 dimers are expressed at the cell surface and in the cytoplasm, can be visualized in non-reducing SDS-PAGE, and are disrupted by DTT. **(A)** Immunoblot of reduced total lysate (“INPUT”) and HA-specific immunoprecipitated (“IP:HA”) fractions from tsA201 cells 48 hours after transient co-expression of 3xHA-Na_v_1.5, 3xFLAG-Na_v_1.5, and 1xGFP-hTpr. Nuclear basket protein, which had a similarly sized plasmid, was used as a transfection control. Immunoprecipitation was performed with HA-coupled magnetic beads. Endogenous α1-syntrophin was used as a positive control for co-immunoprecipitation with Na_v_1.5, and α-actin was a negative control. **(B)** Schematic illustration of the NanoBiT assay, whereby different quantities of untagged Na_v_1.5 are co-expressed together with the luminesce-producing pair of Na_v_1.5-LgBiT and Na_v_1.5-SmBiT. Background signal was determined by co-expression of Na_v_1.5-LgBiT with a non-interacting control, HaloTag-SmBiT. **(C)** In living cells, the presence of untagged Na_v_1.5 significantly decreased the luminescence produced by the interaction between Na_v_1.5-LgBiT and Na_v_1.5-SmBiT. This signifies that untagged Na_v_1.5 competed for the interaction between Na_v_1.5-LgBiT and Na_v_1.5-SmBiT, therefore the observed luminescence signal was Na_v_1.5-specific. Results of the NanoBiT assay are presented as the relative intensity of luminescence (indicating the level of protein-protein interactions) normalized to the fluorescence (indicating the number of cells) of tsA201 cells 48 hours after transfection. Each dataset was normalized to “+ empty vector” control. Data are presented as mean ± SEM from three biological replicates. Multiplicity-adjusted *p*-values, calculated with one-way ANOVA and post hoc Tukey’s multiple comparisons, are indicated in each panel. **(D)** Immunoblot of the non-reduced total lysate from tsA201 cells 48 hours after transient expression of untagged Na_v_1.5 alone and with 1xGFP-Na_v_1.5. White triangles indicate bands corresponding to monomeric untagged Na_v_1.5 and monomeric 1xGFP-Na_v_1.5. The difference in monomer sizes is affected by the GFP tag (∼30 kDa). Thus, the upper bands (∼460 kDa) revealed with anti-Na_v_1.5 and anti-GFP antibodies were considered to indicate dimers of Na_v_1.5. **(E)** Immunoblots of native Na_v_1.5 obtained from lysates of excised mouse hearts prepared using a benchtop homogenizer, direct lysis of isolated cardiomyocytes, and a tissue homogenizer. **(F)** Immunoblot of non-reduced and reduced total lysates of tsA201 cells 48 hours after transient co-expression of mouse and human Na_v_1.5. Dimers and monomers of Na_v_1.5 were revealed with an anti-Na_v_1.5 antibody. Endogenous calnexin was used as a loading control.

To resolve Na_v_1.5 dimers at the expected molecular weight of ∼460 kDa, we performed non-reducing 4%–8% gradient SDS-PAGE. To deduce whether the observed high molecular weight band represented the Na_v_1.5-Na_v_1.5 complex rather than an aspecific band recognized by the antibody used during immunoblotting, we co-expressed non-tagged Na_v_1.5 with green fluorescent protein (GFP)-tagged Na_v_1.5. Accordingly, a ∼460-kDa band was revealed with both antibodies, against Na_v_1.5 and against GFP, while differences in the sizes of corresponding monomeric Na_v_1.5 proteins (∼30 kDa for GFP tag) confirmed the validity of the constructs used (Fig. 1D). Therefore, we concluded that the ∼460-kDa band represented the dimer of Na_v_1.5 proteins, which could also be represented by the SDS-resistant aggregate.

Because our non-reducing SDS-PAGE yielded robust results for detecting the putative Na_v_1.5 dimer/aggregated, we decided to apply this method to native tissues. Using a C57BL/6 mouse model with two untagged wild-type alleles for *SCN5A*, we performed three different types of protein extraction on excised hearts: tissue homogenizer, benchtop homogenizer, and direct lysis of isolated cardiomyocytes. Importantly, regardless of the protein extraction method used, only monomers of Na_v_1.5 were detected (Fig. 1E). To verify that our result was not caused by species specificity, we carried out non-reducing SDS-PAGE on protein lysates from tsA201 cells in parallel expressing mouse and human Na_v_1.5 proteins (Fig. 1F). Immunoblotting analysis revealed the Na_v_1.5-specific bands corresponding to the expected size of its monomeric state (between 171-kDa and 238-kDa of protein standard bands) as well as its putative dimeric/aggregate state (∼460-kDa of protein standard band) observed in Fig. 1D (Fig. 1F). Interestingly, application of a reducing agent, 100 mM DTT, decreased the intensity of putative Na_v_1.5 dimer/aggregate band and increased the amount of Na_v_1.5 monomers (Fig. 1F). This could be due to the disruption of Na_v_1.5 dimers into monomers, suggesting the involvement of covalent disulfide bridges. Therefore, we concluded that these putative Na_v_1.5 dimers/aggregates were at least partially sensitive to the reducing agent and were absent in non-diseased mouse cardiomyocytes.

### Nav1.5 dimers are present at the plasma membrane, but are not primarily formed by disulfide bonds

Functional Na_v_1.5 proteins are typically targeted to the plasma membrane. To assess whether the interacting α-subunits were trapped intracellularly, we performed a biotinylation assay. Our results suggest that both monomers and putative dimers/aggregates of Na_v_1.5 were present at the cell surface when heterologously expressed in tsA201 (Fig. 2A).

**Figure 2.**
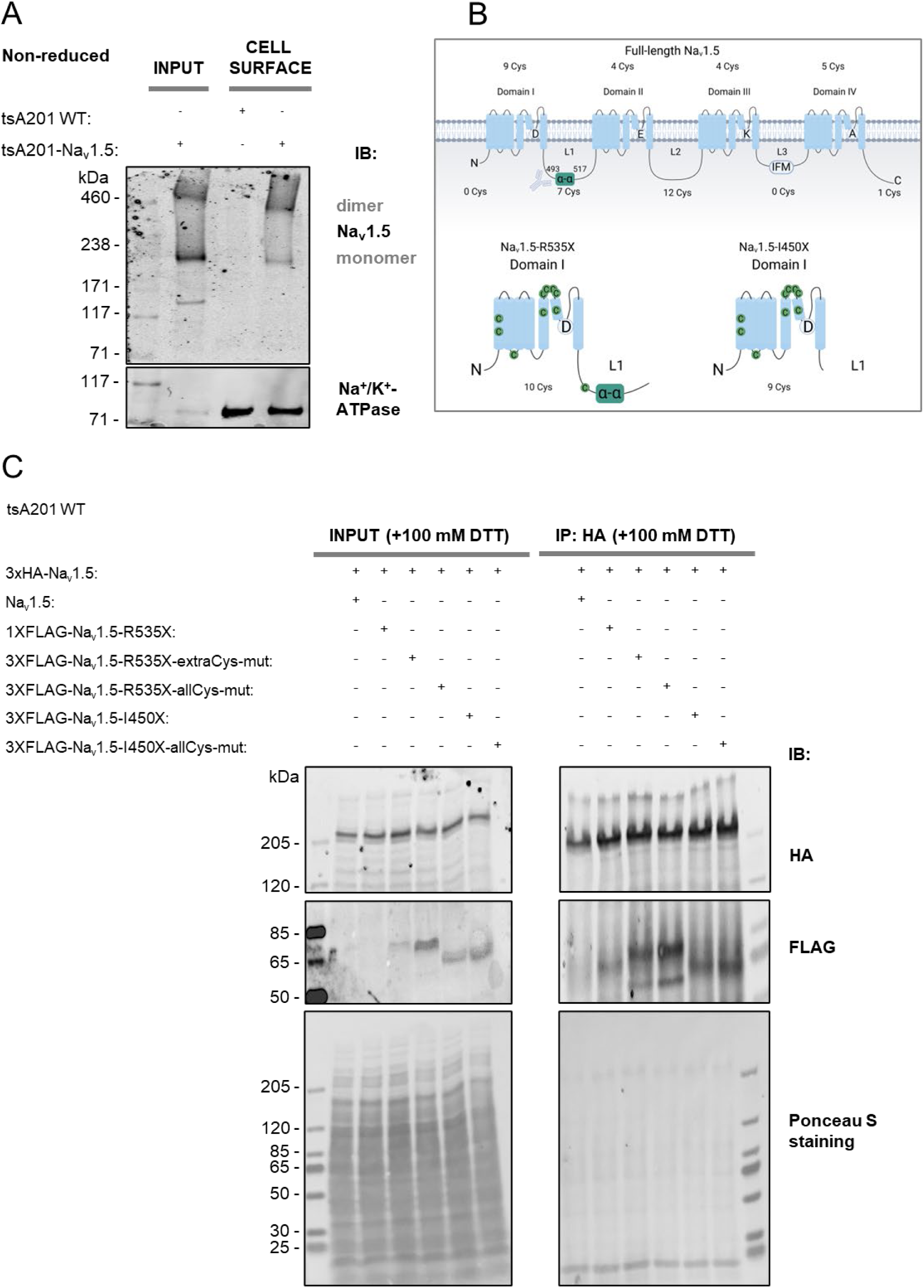
Disulfide bridges are involved in oligomerization of Na_v_1.5 proteins but are not exclusive to this process. (A) Immunoblot of non-reduced total lysate (“INPUT”) and biotinylated (“CELL SURFACE”) fractions of wild-type (WT) and tsA201 cells stably expressing Na_v_1.5. Dimers and monomers of Na_v_1.5 were revealed with an anti-Nav1.5 antibody. Endogenous α1-Na^+^/K^+^-ATPase was used as a positive control for the biotinylated fraction. (B) Schematic illustration of full-length Na_v_1.5 with numbers of Cys corresponding to each domain indicated, and truncated Na_v_1.5 constructs with total numbers of Cys. Na_v_1.5-R535X comprises the presumable α-α-subunit interaction site between Arg493-Arg517, while Na_v_1.5-I450X does not include this site. (C) Immunoblot of the reduced total lysate (“INPUT”) and HA-specific immunoprecipitated (“IP:HA”) fractions of tsA201 cells 48 hours after transient co-expression of 3xHA-Na_v_1.5, untagged Na_v_1.5, 1xFLAG-Na_v_1.5-R535X, 3xFLAG-Na_v_1.5-R535X-extraCys-mut, 3xFLAG-Na_v_1.5-R535X-allCys-mut, 3xFLAG-Na_v_1.5-I450X, and 3xFLAG-Na_v_1.5-I450X-allCys-mut. Ponceaus S staining was used as a loading control.

Since Cys bridges are typically extracellular, this finding was in line with our previous suggestion that disulfide bridges may play a role in linking two Na_v_1.5 proteins together. However, considering that α-α-subunit interactions could also be represented by the aggregated misfolded proteins, we also assessed the possibility of non-native disulfide bridges formed via intracellular Cys.

The full-length Na_v_1.5 (UniProtKB: Q14524-1) has 42 Cys residues in total (Fig. 2B). Importantly, a previous study reported on the existence of a direct α-α-subunit interaction site between Arg493-Arg517 (Fig. 2B) (Clatot et al., 2017). Like that study, we used the truncated Na_v_1.5 proteins, Na_v_1.5-Arg535X and Na_v_1.5-Ile450X, to additionally investigate the role of Arg493-Arg517 for the direct interaction (Fig. 2B). Moreover, to identify which Cys could be responsible for the intermolecular disulfide bonds between Na_v_1.5 proteins, we mutated Cys into Ala. For this, we created Na_v_1.5-Arg535X constructs with only 4 extracellular Cys (Na_v_1.5-Arg535X-extraCys-mut) and all 10 Cys being mutated (Na_v_1.5-Arg535X-allCys-mut). Similarly, we mutated all 9 Cys of Na_v_1.5-Ile450X (Na_v_1.5-Ile450X-allCys-mut). Co-immunoprecipitation analysis between full-length Na_v_1.5 and the aforementioned truncated Na_v_1.5 proteins revealed that mutating only 4 extracellular Cys or all 10 Cys was insufficient to abolish the ability of Na_v_1.5-Arg535X protein to interact with the full-length Na_v_1.5 (Fig. 2C). Furthermore, Na_v_1.5-Ile450X, lacking the presumed α-α-subunit interaction site between Arg493-Arg517, also preserved its binding with the full-length Na_v_1.5, and this binding was not affected by the absence of Cys (Fig. 2C). Therefore, we concluded that Arg493-Arg517 was not responsible for linking two Na_v_1.5 proteins. Moreover, although Na_v_1.5 multimers/aggregates could have been at least partially supported via disulfide bridges, as observed in Fig. 1F, they were not the main cause of its oligomerization/aggregation.

### Nav1.5 dimers are unaffected by co-expression with Navβ-subunits

Na_v_β-subunits, particularly Na_v_β2 and Na_v_β4, are known to form intermolecular disulfide bridges within themselves as well as with α-subunits of VGSCs [reviewed in (Iamshanova et al., 2023)]. Notably, a previous study proposed that α-α-subunit interactions of Na_v_1.5 were regulated by the presence of Na_v_β1 (Mercier et al., 2012). Although we and other groups reported the self-interactions of Na_v_1.5 proteins in the absence of Na_v_β-subunits (Iamshanova et al., 2024; Clatot et al., 2017), we decided to revise this question in this study. First, we compared the ratios of Na_v_1.5 dimers/monomers with and without individual co-expression of Na_v_β1B, Na_v_β1, Na_v_β2, Na_v_β3, and Na_v_β4 subunits (Fig. 3A, C). Second, we analyzed the amount of haemagglutinin (HA)-tagged Na_v_1.5 that co-immunoprecipitated with FLAG-Na_v_1.5 in the absence and presence of Na_v_β-subunits co-expression (Fig. 3B, D). Third, we estimated levels of the putative Na_v_1.5 dimers between the aforementioned conditions in living cells (Fig. 3E). Because we did not observe an effect in any experimental procedure, we concluded that Na_v_β-subunits are indeed dispensable for the homo-oligomerization/aggregation of Na_v_1.5 in our heterologous expression system.

**Figure 3.**
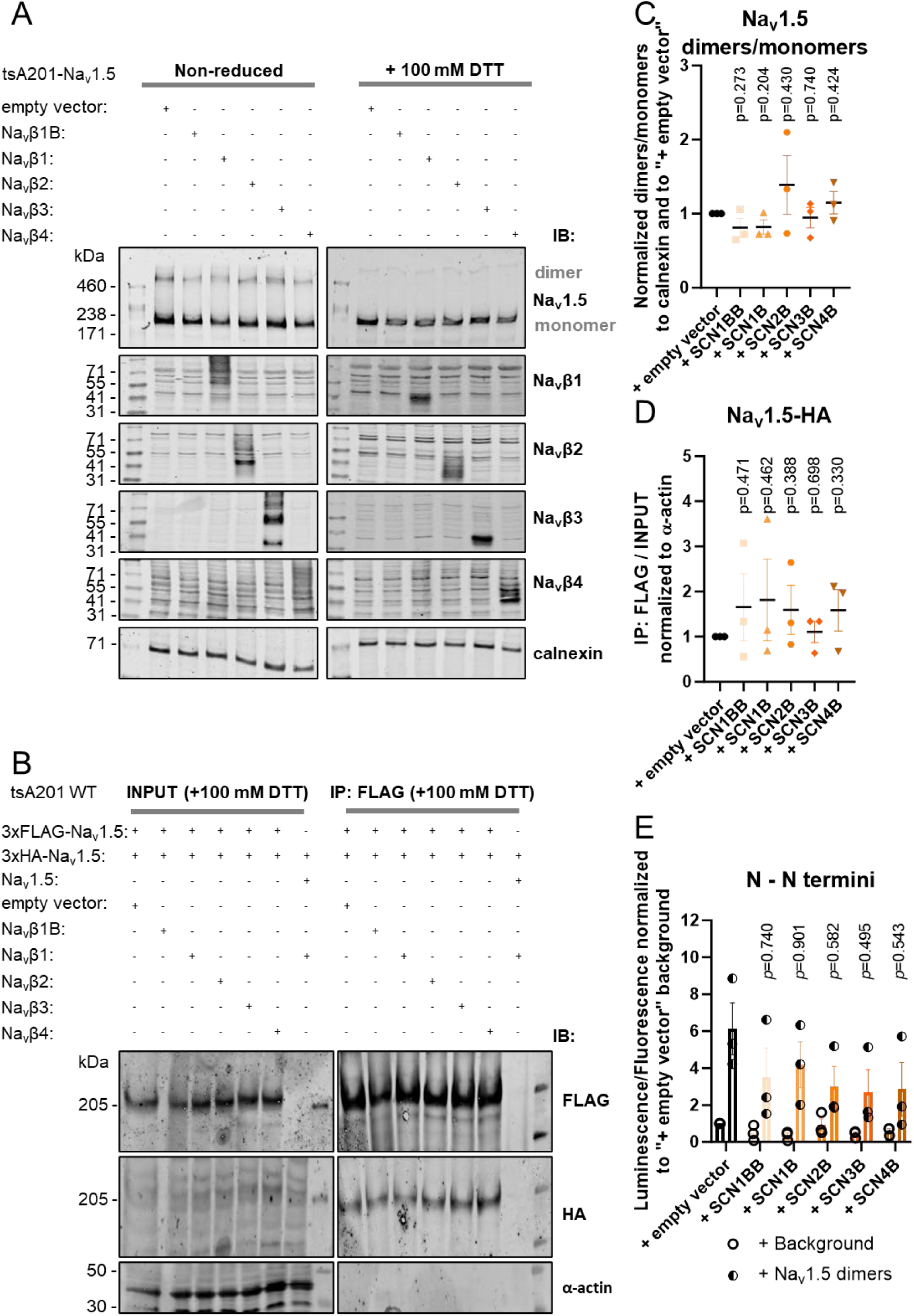
Na_v_1.5 dimers are unaffected by co-expression of Na_v_β-subunits. **(A)** Immunoblot of non-reduced and reduced total lysate fractions of tsA201 cells stably expressing Na_v_1.5 after 48 hours of transient expression of Na_v_β1B, Na_v_β1, Na_v_β2, Na_v_β3, and Na_v_β4. Dimers and monomers of Na_v_1.5 were revealed with anti-Na_v_1.5. Endogenous calnexin was used as a loading control. Of note, Na_v_β1B is not expected to have an epitope for detection with anti-Na_v_β1. **(B)** Immunoblot of the reduced total lysate (“INPUT”) and FLAG-specific immunoprecipitated (“IP:FLAG”) fractions of tsA201 cells 48 hours after transient co-expression of 3xHA-Na_v_1.5 and 3xFLAG-Na_v_1.5 with and without Navβ1B, Navβ1, Navβ2, Navβ3, and Navβ4. Endogenous α-actin was used as a negative control. **(C)** Intensities of Na_v_1.5 dimers/monomers from (A) were normalized to calnexin and the control condition (“+ empty vector”). Data are presented as mean ± SEM from three biological replicates. Individual *p*-values, calculated with a one-sample two-tailed t-test with hypothetical mean value = 1, are indicated in each panel. **(D)** Intensities of Na_v_1.5-HA in (B) from “IP:FLAG” were normalized to their corresponding intensities from INPUT and further normalized to calnexin and the control condition (“+ empty vector”). Data are presented as mean ± SEM from three biological replicates. Individual *p*-values, calculated with a one-sample two-tailed t-test with hypothetical mean value = 1, are indicated in each panel. **(E)** Results of the NanoBiT assay are presented as the relative intensity of luminescence (indicating the level of protein-protein interactions) normalized to fluorescence (indicating the number of cells) of tsA201 cells 48 hours after transfection. Each dataset was normalized to “+ empty vector” control. Data are presented as mean ± SEM from three biological replicates. The multiplicity-adjusted *p*-values, calculated with one-way ANOVA and post hoc Tukey’s multiple comparisons, are indicated in each panel.

### Transmembrane domains strongly interact with full-length Nav1.5 and reduce its protein level

Because our data did not confirm the exclusivity of the previously described Arg493-Arg517 site for α-α-subunit interactions of Na_v_1.5 (Fig. 2C) (Clatot et al., 2017), we decided to revise the interaction sites by co-immunoprecipitation analysis between different parts of Na_v_1.5 and the full-length protein (Fig. 4A). Strikingly, the presence of transmembrane domains I–IV decreased total levels (both putative dimers/aggregates and monomers) of full-length Na_v_1.5 (Fig. 4A, B). This could be explained by increased aspecific hydrophobic interactions between the large transmembrane domains of Na_v_1.5 that would lead to protein aggregation and activation of UPR with subsequent proteasomal degradation (Fig. 4C). Indeed, although the total protein quantity was diminished, the interaction between transmembrane domains I–IV and full-length Na_v_1.5 remained very strong (Fig. 4B). Furthermore, we detected potent interactions between full-length Na_v_1.5, its intracellular N-terminal domain (NTD), and the linker between domains I and II (named L1) (Fig. 4B). Overall, these results indicate that the heterologusly expressed Na_v_1.5 proteins interact with each other not due to a specific amino acid sequence but rather within multiple regions.

**Figure 4.**
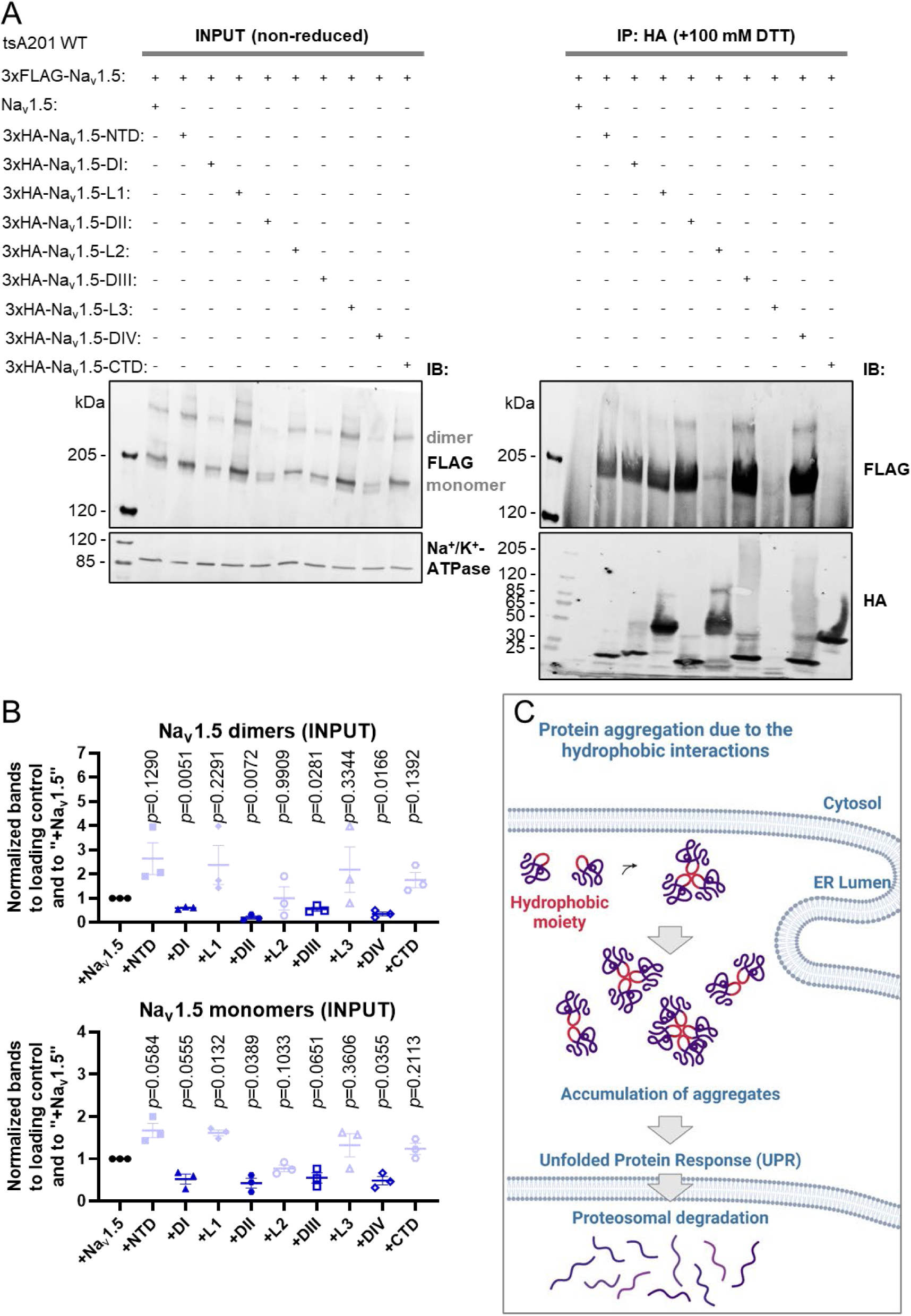
α-α-Subunit interactions occur between full-length Na_v_1.5 and all four transmembrane domains, the N-terminal domain, and the linker between domains I and II. (A) Immunoblot of non-reduced total lysate (“INPUT”) and reduced HA-specific immunoprecipitated (“IP:HA”) fractions of tsA201 cells 48 hours after transient co-expression of 3xFLAG-Na_v_1.5 with untagged Na_v_1.5, 3xHA-Na_v_1.5-NTD, 3xHA-Na_v_1.5-DI, 3xHA-Na_v_1.5-L1, 3xHA-Na_v_1.5-DII, 3xHA-Na_v_1.5-L2, 3xHA-Na_v_1.5-DIII, 3xHA-Na_v_1.5-L3, 3xHA-Na_v_1.5-DIV, and 3xHA-Na_v_1.5-CTD. In total lysate, Na_v_1.5 dimers and monomers were revealed with an anti-FLAG antibody. The immunoprecipitated domains of Na_v_1.5 were revealed with an anti-HA antibody, while co-immunoprecipitated full-length Na_v_1.5 proteins were revealed with an anti-FLAG antibody. Endogenous α1-Na^+^/K^+^-ATPase was used as a loading control for the total lysate fraction. (B) Intensities of non-reduced Na_v_1.5 dimers and non-reduced Na_v_1.5 monomers from (A) were normalized to α1-Na^+^/K^+^-ATPase and the condition with co-expression of untagged full-length Na_v_1.5 (“+Na_v_1.5”). Data are presented as mean ± SEM from three biological replicates. Individual *p*-values, calculated with a one-sample two-tailed t-test with hypothetical mean value = 1, are indicated in each panel. (C) Schematic illustration of how hydrophobic interactions between transmembrane domains of Na_v_1.5 might promote protein aggregation and UPR.

### Nav1.5 dimerization levels are regulated by its protein expression level and core N-linked glycosylation in the ER

All α-α-subunit interactions of VGSCs were previously shown in a heterologous overexpression system. In this case, there is a possibility that the overload and/or unsuitability of protein folding machinery could lead to the aggregation of ectopic proteins (Fig. 5A). Thus, we decided to assess the oligomerization/aggregation of Na_v_1.5 while controlling the level of its overexpression. To achieve this, we compared the ratios of the putative reducing agent-sensitive Na_v_1.5 dimers/aggregates to its monomers produced under promoters with different expression strengths (Fig. 5B). According to our results, the Na_v_1.5 dimer/monomer ratio was directly proportional to the level of heterologous protein expression (Fig. 5C). Interestingly, the strongest promoter, cytomegalovirus (CMV), also led to the production of monomeric Na_v_1.5 of a smaller size compared with monomeric Na_v_1.5 proteins produced with SV40 and Ubiquitin C (UbC) promoters (Fig. 5B). This encouraged us to further investigate whether a slight change in the size of monomeric Na_v_1.5 under strong levels of expression was associated with the insufficiency of post-translational modifications resulting from saturated protein folding machinery (Fig. 5A). One of the first post-translational modifications is N-glycosylation, which occurs while the nascent peptide is being synthesized to assist with correct protein folding in the ER (Fig. 5D). Both native and heterologous Na_v_1.5 were shown to be N-glycosylated (Mercier et al., 2015). Therefore, we used an inhibitor of N-core glycosylation, tunicamycin (TUN), to test its importance for the heterologous oligomerization/aggregation of Na_v_1.5 (Fig. 5D). TUN prevents the addition of oligosaccharides to nascent polypeptides, leading to their misfolding and accumulation in the ER (Fig. 5D). This, in turn, induces ER stress followed by activation of the UPR and proteasomal degradation (Fig. 5D). To constrain the latter and preserve Na_v_1.5 dimers/aggregates possibly evoked due to the affected N-glycosylation, we used the cell-permeable proteasomal inhibitor MG132 (Fig. 5D). Because the presence of TUN and MG132 might significantly alter the total amount of Na_v_1.5 protein, we compared the differences between Na_v_1.5 dimer/monomer ratios under various conditions by normalizing them to treatment with the highest total protein amount, i.e. MG132 alone. Accordingly, our results confirmed that TUN alone induced the degradation of heterologous protein and reduced the size of the band corresponding to monomeric Na_v_1.5, whereas the addition of MG132 rescued the amount of heterologous Na_v_1.5 (Fig. 5E, F). Furthermore, the amount of non-reduced Na_v_1.5 monomers decreased under the condition of combined TUN and MG132 treatment, while the reduced Na_v_1.5 monomers tended to increase (Fig. 5E, F). This result could signify that the combination of TUN with MG132 facilitated the accumulation of Na_v_1.5 oligomers compared with MG132 treatment alone. Altogether, our results suggest that a high production rate of heterologous Na_v_1.5 might saturate the ER folding machinery, and this, in turn, as observed with impaired N-glycosylation, could promote its dimerization/aggregation.

**Figure 5.**
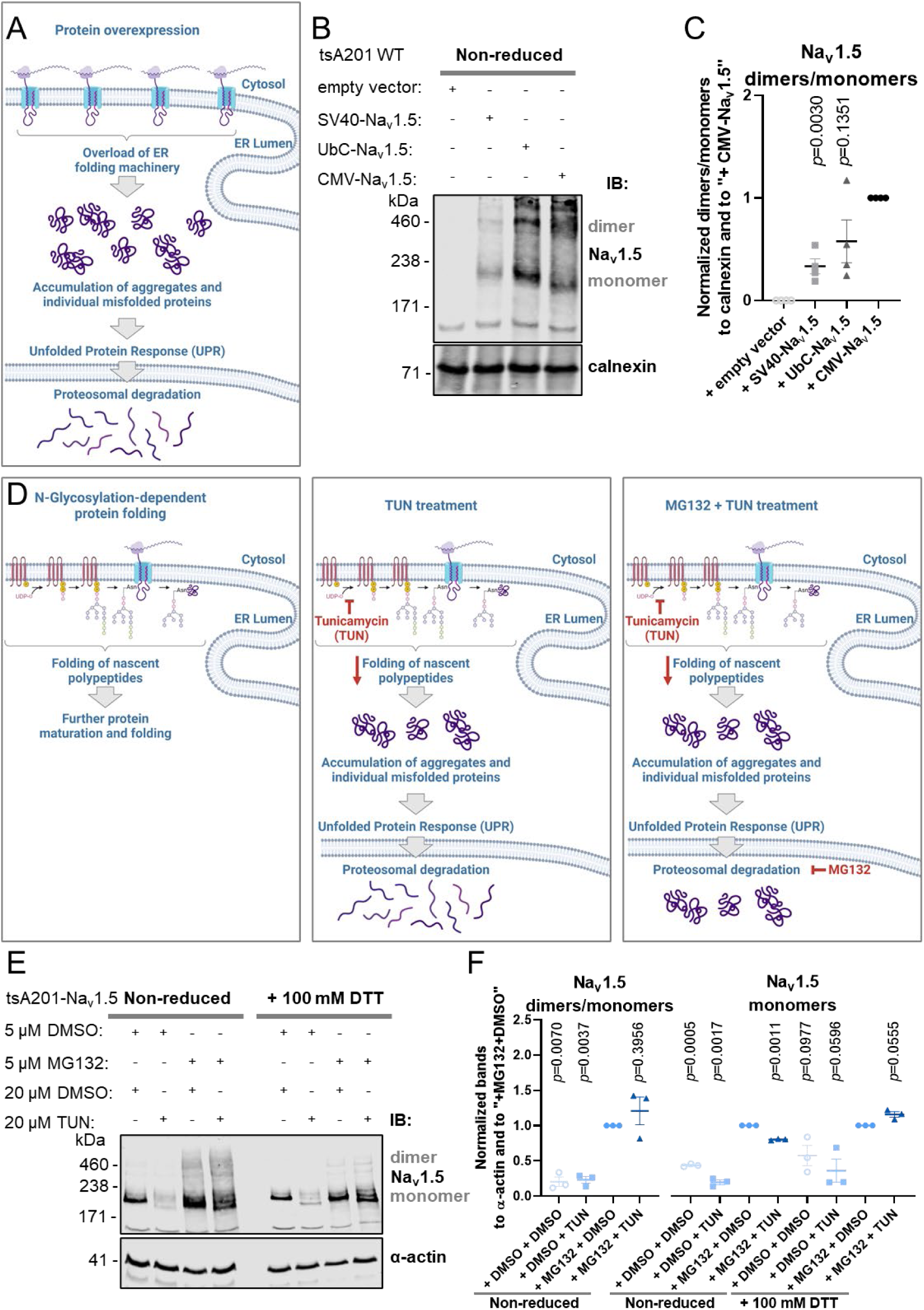
Na_v_1.5 dimerization is increased at higher expression levels and with ER stress induction. **(A)** Schematic illustration of how protein overexpression might overload ER folding machinery and trigger UPR. **(B)** Immunoblot of the non-reduced total lysate fraction of tsA201 cells after 48 hours of transient expression of Na_v_1.5 with promoters of different strengths. Dimers and monomers of Na_v_1.5 were revealed with anti-Na_v_1.5. Endogenous calnexin was used as a loading control. **(C)** Intensities of Na_v_1.5 dimers/monomers from (B) were normalized to calnexin and the condition with the strongest promoter (“+CMV-Na_v_1.5”). Data are presented as mean ± SEM from three biological replicates. Individual *p*-values, calculated with a one-sample two-tailed t-test with hypothetical mean value = 1, are indicated in each panel. **(D)** Schematic illustration of how TUN treatment (prevents N-core glycosylation) and MG132 treatment (inhibits proteasomal degradation) affect protein folding and UPR. **(E)** Immunoblot of non-reduced and reduced total lysate fractions of tsA201 cells stably expressing Na_v_1.5 with the indicated combinations of TUN and MG132 treatments. Endogenous α-actin was used as a loading control. **(F)** Intensities of non-reduced Na_v_1.5 dimers/monomers, as well as non-reduced and reduced Na_v_1.5 monomers from (B), were normalized to α-actin and the condition with inhibited proteasomal degradation (“+MG132+DMSO”). Data are presented as mean ± SEM from three biological replicates. Individual *p*-values, calculated with a one-sample two-tailed t-test with hypothetical mean value = 1, are indicated in each panel.

## Discussion

By using non-reducing SDS-PAGE, we were able to resolve putative Na_v_1.5 dimers in the heterologous expression system represented by tsA201 cells, but not in native mouse cardiomyocytes. These putative Na_v_1.5 dimers were shown to be partially sensitive to the reducing agent, depending on the ER stress and the presence of transmembrane domains. This could signify that the apparent dimers of Na_v_1.5 are likely being represented by the SDS-resistant aggregates, supported by the non-native disulfide bonds.

We show that these dimers/aggregates were unaffected by Na_v_β-subunits but rather relied on the level of *SCN5A* overexpression. We identified that strong interactions occurred between the full-length Na_v_1.5 and its NTD, DI, L1, DII, DIII, and DIV in our cell model.

Our study validates the occurrence of dimerized human VGSC α-subunits, formerly reported by us and other groups (Mercier et al., 2012; Clatot et al., 2012, 2017; Salvage et al., 2022; Rühlmann et al., 2020; Iamshanova et al., 2024; Salvage et al., 2020). In addition to the previously described homodimerization site of Na_v_1.5 at L1 (Clatot et al., 2017), our group previously observed close proximity between the full-length protein and its truncated NTD (Wang et al., 2020). In the present study, we confirm direct interactions between full-length Na_v_1.5 and its NTD, as well as L1. Recently, another inter-channel dimerization of a VGSC α-subunit was reported and suggested to take place through its voltage-sensing domain (Sumino et al., 2023). In support of these observations, our results show that all four transmembrane domains (i.e., DI, DII, DIII, and DIV) were indeed able to strongly interact with full-length Na_v_1.5.

Interestingly, two-distinct pools of *N*-glycosylated Na_v_1.5 channels were reported for the human embryonic kidney cell line HEK293T (RRID:CVCL_0063), a heterologous expression system highly similar to ours (Mercier et al., 2015). A previous study proposed the co-existence of fully mature Na_v_1.5 channels that underwent post-*cis*-Golgi glycosylation together with partially mature *N*-glycan-restricted Na_v_1.5 proteins (Mercier et al., 2015). Even though the latter did not undergo terminal glycosylation, both Na_v_1.5 pools were shown to traffic to the plasma membrane (Mercier et al., 2015). The authors suggested that the unconventional Golgi-independent pathway of membrane transport could be potentially used for the clearance of Na_v_1.5 proteins accumulating in the ER to prevent ER stress (Mercier et al., 2015). Similarly, we suggest a possible link between Na_v_1.5 dimerization/aggregation and its post-translational modifications, such as *N*-glycosylation. According to our results, preventing early *N*-glycosylation of nascent polypeptides with TUN while blocking proteasomal degradation with MG132 tended to promote Na_v_1.5 dimerization/aggregation. We also show that Na_v_1.5 dimers/aggregates reached the plasma membrane. In line with these findings, Förster resonance energy transfer studies demonstrated that Na_v_1.5 α-subunits interacted at the cell surface and that these interactions occurred before protein trafficking to the plasma membrane (Clatot et al., 2017).

Previously, Na_v_β1 was reported to mediate α-α-subunit interactions between Na_v_1.5 proteins (Mercier et al., 2012). However, our current data demonstrate that dimerization/aggregation of Na_v_1.5 was unaffected by the presence of Na_v_β1, its alternative splicing form Na_v_β1B, or subunits Na_v_β2, Na_v_β3, and Na_v_β4. This was not surprising to us, as other studies showing oligomerization of human VGSC α-subunits did not use co-expression of Na_v_β-subunits (Clatot 2012, Clatot 2017, Rhulmann 2020). In addition, another study concluded that clusterization of Na_v_1.5 is possible with and without Na_v_β3 (Salvage et al., 2020). Unexpectedly, the α-α-subunit interaction site previously described for Na_v_1.5 as Arg493-Arg517 region at L1 was not proven to be exclusive in our hands (Clatot et al., 2017). Previously, the authors demonstrated that truncated GFP-tagged constructs Na_v_1.5-Ile450X and Na_v_1.5-Asn470X were unable to co-immunoprecipitate with full-length HA-tagged Na_v_1.5 (Clatot et al., 2017). However, according to our results, Na_v_1.5-Ile450X strongly interacted with full-length Na_v_1.5. Indeed, such an interaction could be mediated via NTD and DI, parts of Na_v_1.5-Ile450X that by themselves were shown to strongly bind to full-length Na_v_1.5 in our current work.

Another discrepancy with our study comes from the previously shown migration of human Na_v_1.5 and Na_v_1.7 as monomers in both reduced and non-reduced states (Rühlmann et al., 2020). Based on these observations, the authors argued against the existence of disulfide bonds between α-subunits but not against non-covalent bonds (Rühlmann et al., 2020). However, our results indicate that Na_v_1.5 dimers/aggregates were reducing agent-dependent, even though disulfide bonds did not appear to be the primary cause of α-α-subunit interactions. One possible explanation could be technical: we used Tris-acetate SDS-PAGE in contrast to SDS-urea-PAGE used in previous work. As a strong protein denaturant, urea might assist in protein unfolding and/or stabilization of the unfolded state. If this is the case and non-native disulfide bridges appear during partial protein refolding as a result of our experimental procedures, then it would not explain the absence of Na_v_1.5 dimers/aggregates in native cardiac tissues/cells when using the same non-reducing Tris-acetate SDS-PAGE protocol. Interestingly, for the 30 proteins analyzed, no correlation was found between the extent of urea-induced protein unfolding and the presence/absence of disulfide bridges in the native form (Candotti et al., 2013). Additionally, another explanation could be experimental: cellular machinery providing for post-translational modifications, including glycosylation, is significantly different between the heterologous expression system used in this study, human tsA201 cells, and *Xenopus laevis* oocytes used in previous work (Rühlmann et al., 2020).

### Strengths/limitations

Importantly, most of our data were obtained in a heterologous expression system using a protein overexpression approach. Thus, it is possible that the observed interactions and effects on Na_v_1.5 dimerization/aggregation were largely affected by non-native ectopic gene expression.

Another limitation of our study is that by using non-reducing Tris-acetate SDS-PAGE, we were only able to resolve putative Na_v_1.5 dimers that primarily depended on the disulfide bonds. Therefore, whenever possible, we performed co-immunoprecipitation analysis as an additional confirmation of the α-α-subunit interactions.

### Implications

Biochemical interactions between α-α-subunits of Na_v_1.5 were largely used to describe/explain the dominant-negative effect (DNE) of *SCN5A* (reviewed in 10). DNE was shown for both native and heterologous cell models (Doisne et al., 2021; O’Neill et al., 2022; Keller et al., 2005). Accordingly, it was hypothesized that DNE occurred due to direct interactions between wild-type and mutant Na_v_1.5 because the trafficking, turnover, and function of the interacting pair would be largely affected (Sottas and Abriel, 2016). However, a direct link between DNE and Na_v_1.5 has yet to be shown, as disruption of the protein-protein interaction between mutant and wild-type channels should lead to the rescue of *I*_Na_.

Our data suggest a link between Na_v_1.5 dimerization/aggregation and ER stress. Although we did not detect Na_v_1.5 dimers in non-diseased mouse cardiomyocytes, such α-α-subunits interactions may occur after ER stress induction. Indeed, fever, pharmacological agents, and age might promote ER stress and are known risk factors for the manifestation of Brugada syndrome, a cardiac disease primarily attributed to *SCN5A* loss-of-function variants (Keller et al., 2005).

## Conclusion

Taken together, our data raise caution regarding challenges related to investigating the oligomerization of Na_v_1.5 in a heterologous expression system. Therefore, further work needs to be carried out, specifically, using an in vivo model where both alleles of *SCN5A* are genetically modified to produce native Na_v_1.5 proteins with different tags.

## Supporting information

Figure S1 with its legend shows analyzed data to complement Figure 1.

## Data availability statement

Data are available in the article itself and its supplementary materials.

## Acknowledgments

Illustrations were created with BioRender.com.

## Author contributions

**O.I, J.S.R.**, and **H.A.** conceptualization. **O.I., A.F.H., S.S., A.S.**, and **S.G.** investigation. **O.I., A.F.H.**, and **S.S.** formal analysis. **O.I., J.S.R.**, and **H.A.** writing – original draft.

## Funding and additional information

This work was funded by the Swiss National Science Foundation [SNF 310030_184783] to H.A.

## Abbreviations

CMV: cytomegalovirus
CTD: C-terminal domain
DNE: dominant-negative effect
DTT: dithiothreitol
HA: haemagglutinin
*I*_na_: Na^+^ current
LDS: lithium dodecyl sulfate
Na^+^: sodium ion
Na_v_1.x: α-subunit of the voltage-gated sodium channel
Na_v_β: β-subunit of the voltage-gated sodium channel
NTD: N-terminal domain
PBS: Phosphate-buffered saline
SDS: PAGE sodium dodecyl sulfate-polyacrylamide gel electrophoresis
TBS: Tris-buffered saline
TUN: tunicamycin
UbC: ubiquitin C
UPR: unfolded protein response
VGSC: voltage-gated sodium channel

---

**Figure S1.**
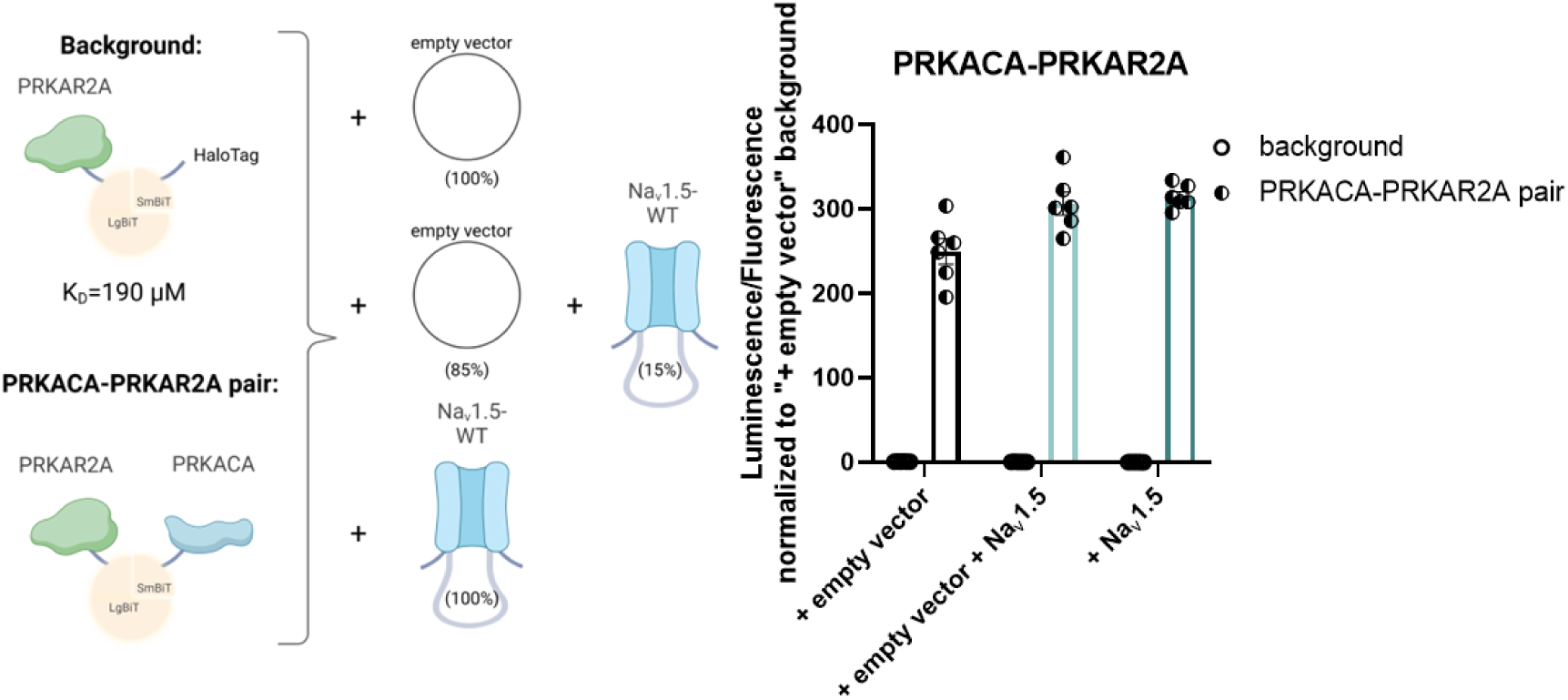
Na_v_1.5 does not affect the interaction between proteins of the control pair represented by PRKACA-PRKAR2A. Schematic illustration of the NanoBiT assay, when co-expressing different quantities of untagged Na_v_1.5 together with the control pair of PRKAR2A-LgBiT and PRKACA-SmBiT. Background signal was determined by co-expression of PRKAR2A-LgBiT with a non-interacting control, HaloTag-SmBiT. Results of the NanoBiT assay are presented as the relative intensity of luminescence (indicating the level of protein-protein interactions) normalized to the fluorescence (indicating the number of cells) of tsA201 cells 48 hours after transfection. Each dataset was normalized to “+ empty vector” control. Data are presented as mean ± SD from six technical replicates.

